## supplemental figures for "The natverse: a versatile computational toolbox to combine and analyse neuroanatomical data"

### Supplementary Information

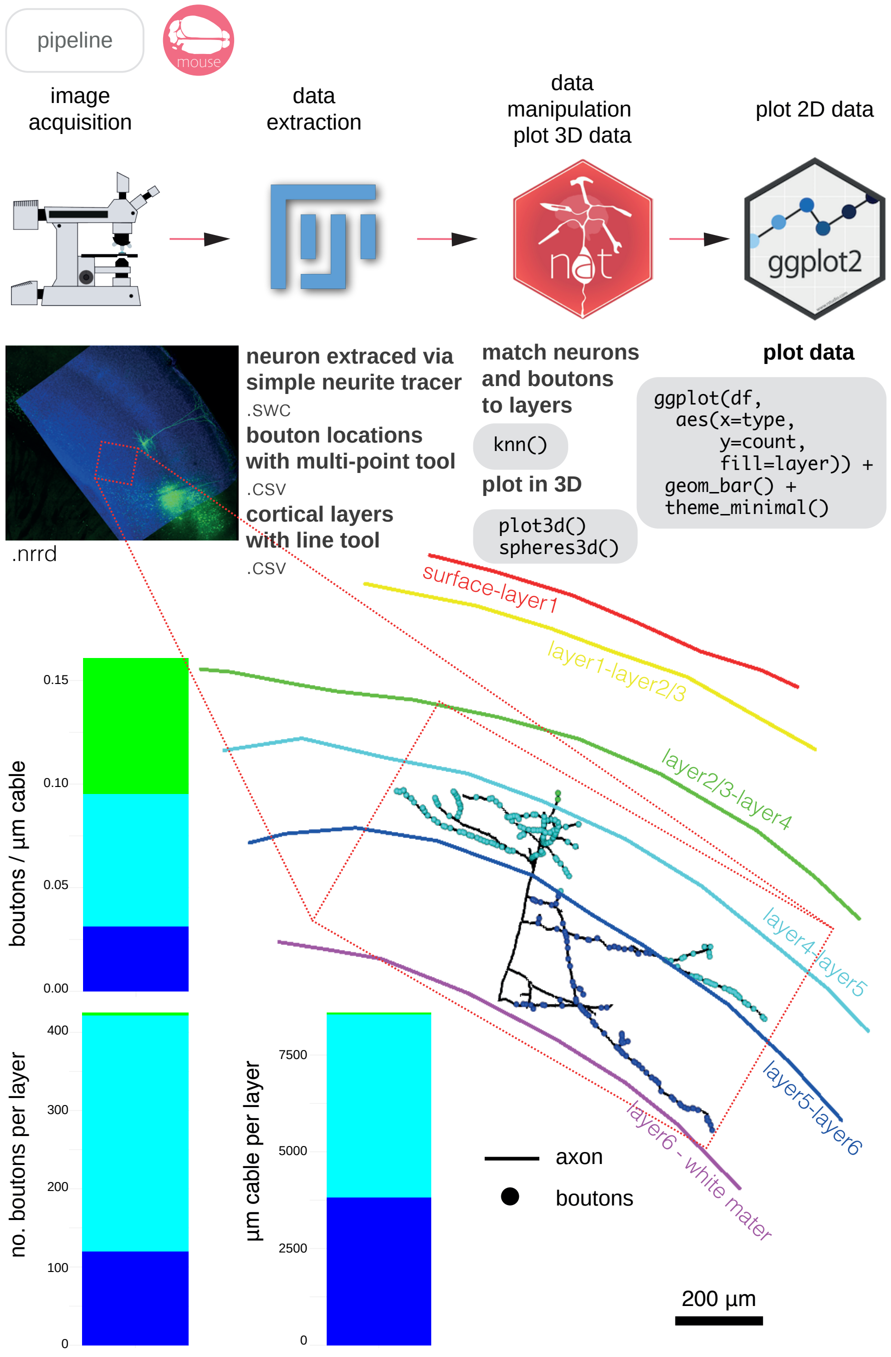


**Supplementary Figure 1:** A simple pipeline for neuron analysis with nat. Bouton placement for a layer 5 pyramidal neuron from the mouse primary somatosensory cortex is examined relative to the neurons' cable length and position within the layered structure of the cortex. Coloured names indicate layer transitions. Data courtesy of A. Vourvoukelis, A. von Klemperer and C.J. Akerman.

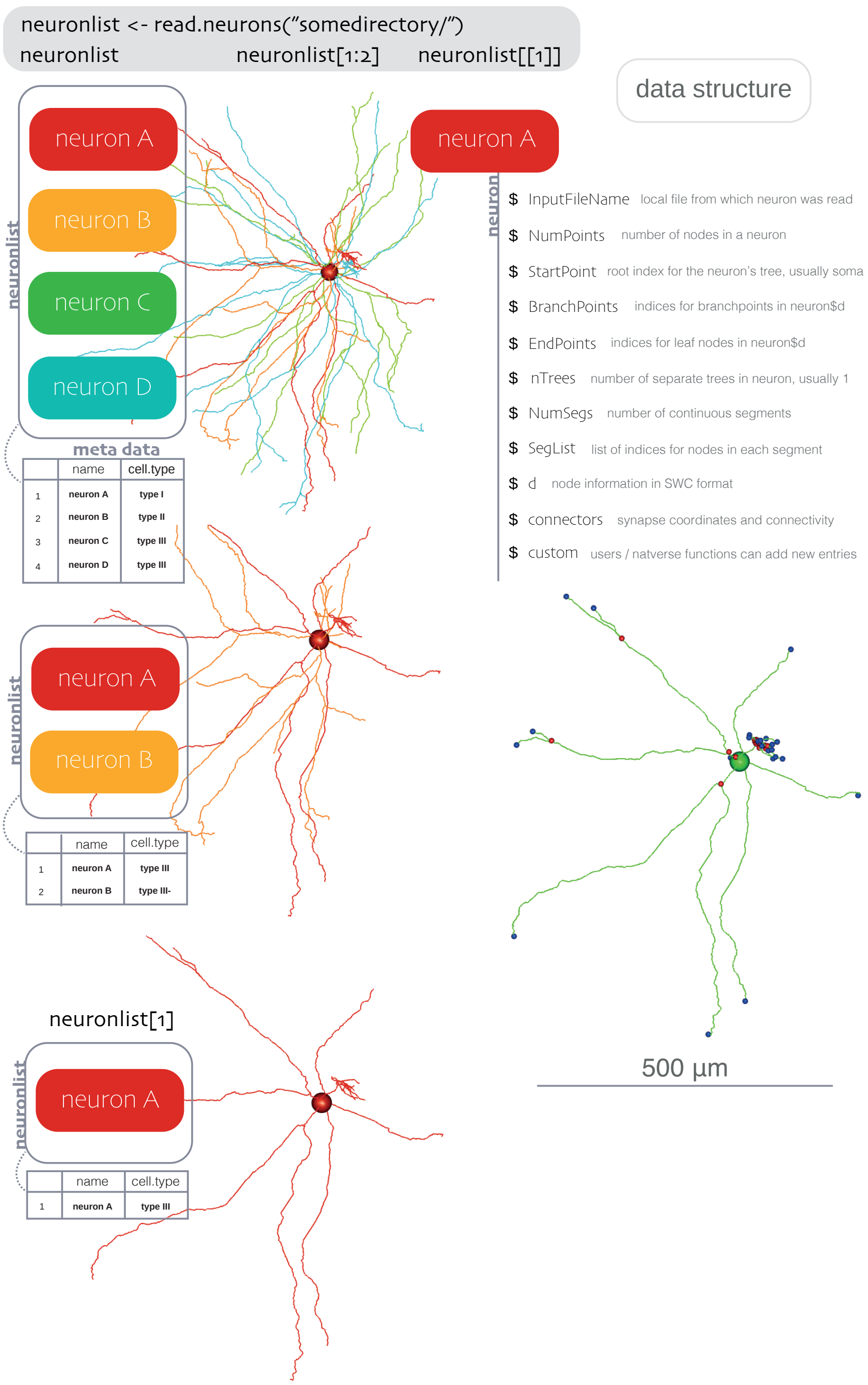


**Supplementary Figure 2:** Schematic representation of the data structure behind `neuron` objects and `neuronlist` objects. Objects of class `neuronlist` are essentially lists of `neuron` objects, representing one or more neurons, with some attached metadata.

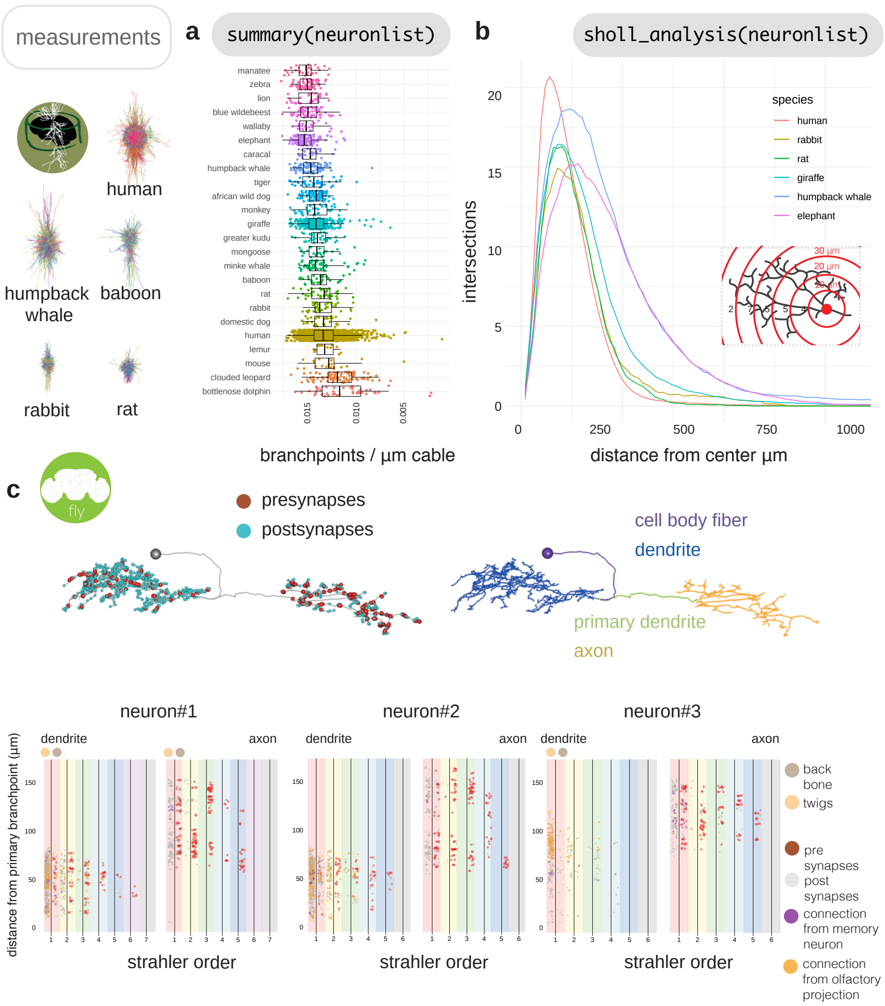


**Supplementary Figure 3: a** We used neuromorphr to obtain information about 40599 mammalian neurons from NeuroMorpho.org, in the brain region ‘neocortex’, from 193 different laboratories. However, it is quite difficult to compare morphologies acquired from different laboratories, that have used different imaging pipelines, neuron reconstruction tools and staining protocols (Farhoodi et al., 2019). We can use neuromorphr to access neuron metadata on NeuroMorpho.org, and choose only those neurons from the same laboratory (Bob Jacobs’), that have been obtained using a Golgi stain and the reconstruction software NeuroLucida, and that are classed as a principal cell (Anderson et al., 2009; Jacobs et al., 2015, 1997; Travis et al., 2005; Jacobs et al., 2016; Anderson et al., 2010; Jacobs et al., 2001, 2018, 2011; Reyes et al., 2016), giving us 3174 neurons. Branch points per micron cable can then be plotted for 24 different species’ cortical principal neurons using ggplot2. **b** Inset, a Sholl analysis (Sholl, 1953) obtains the number of intersections the neurons’ morphologies make with radii of increasing size centered at their roots. Main, the mean intersections for a subset of the species’ cortical principal cells is shown. **c** Right, using synaptic information, it is possible to use a graph theoretic approach which divides the neuron at the point of maximum ‘flow’ having ‘drawn’ a path between each input synapse and each output synapse that pass through every node on the skeleton (Schneider-Mizell et al., 2016). This helps divide a neuron into its dendrites, axon, intervening cable (maximum flow, the primary dendrite) and its cell body fiber (no flow). In insects, the cell body lies outside the neuropil and is connected to its arbour by a single fiber. Left, axon-dendrite split shown for exemplary neuron using `seesplit3d`. **d** The distribution of pre- and post-synapses along three neurons arbors, split by axon and dendrite, then further by Strahler order, then further by presence (backbone) or absence (twigs) of microtubule. Olfactory related neurons (gold), associative memory related neurons (purple) and unknown inputs (grey). In this case, neuron#1 and neuron#2 are of the same cell type and look similar (PD2a1), but neuron#3 differs, indeed it belongs to a different cell type (PD2b1) (Dolan et al., 2018).

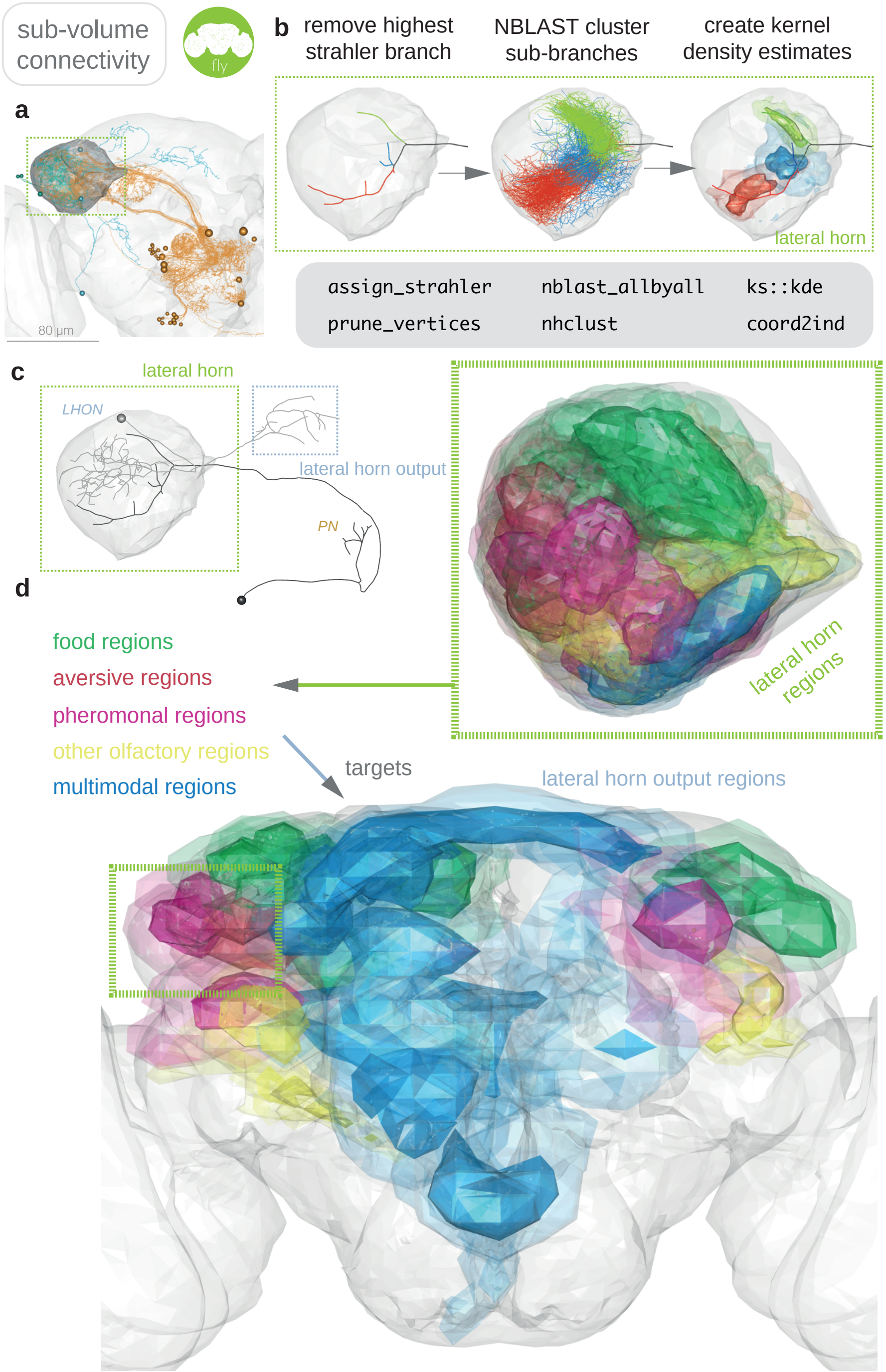


**Supplementary Figure 4: a** The basic connectivity scheme of the lateral horn, a second order olfactory centre in insects. Olfactory projection neurons (in orange) connect to lateral horn neurons (in cyan) (Frechter et al., 2019). **b** To create anatomically meaningful continuous voxels for the lateral horn, rather than random contiguous partitions of our standard neuropil space (Ito et al., 2014), we first removed the highest Strahler order branch (`assign_strahler`) from projection neuron axons' so that their sub-branches could be clustered into 25 separate groups (`nblast`). For each cluster, a 3D weighted kernel density estimate was generated based on 3D points (`xyzmatrix`) extracted from clustered sub-branches, using the R package ks (Duong, 2007). Points spaced on neurites at 1 $\mu$m intervals (`resample`) and weighted as 1 / total number of points in the cluster, so that supervoxels could be directly compared. **c** An 'inclusion' score for each neuron was calculated for each supervoxel by summing the density estimate for each point in the chosen arbor, again sampled at 1 $\mu$m intervals, and normalized by the total number of points in each arbor. An atlas of the lateral horn, colouring supervoxels by the modality/valence of their strongest input neurons. **d** A 'projection' was calculated between each lateral horn voxel and each lateral horn output voxel based on the number of neuronal cell types that have processes in both and the density of this arborisation. An atlas of the lateral horn output regions can then be made, colouring supervoxels by the modality/valence of their strongest input lateral horn supervoxels.

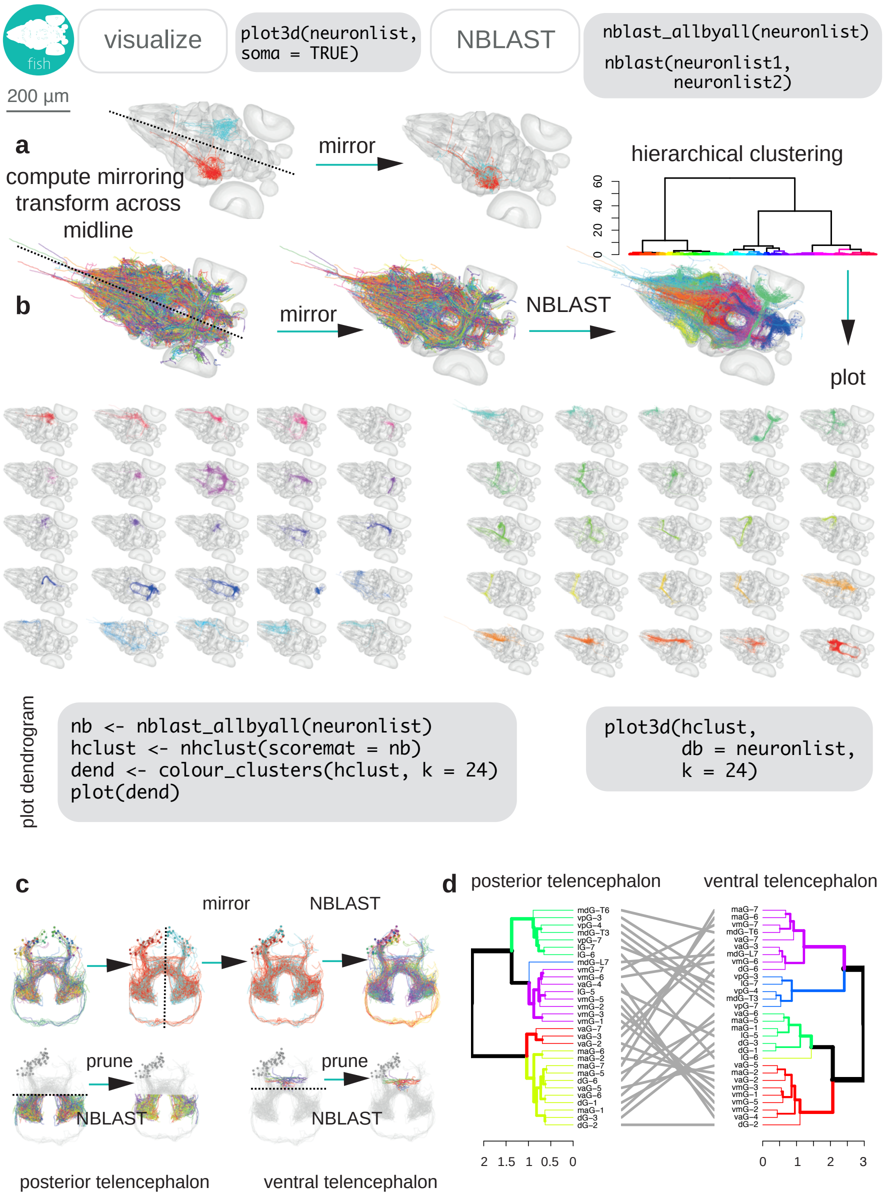


**Supplementary Figure 5: a** Light-level neurons registered to a standard brain for the larval zebrafish (Kunst et al., 2019) can be read into R using fishatlas, and all transformed onto the right hemisphere using the mirroring registration generated by Kunst et al. **b** They can then be NBLAST-ed to discover new cell types. 25 NBLAST clusters are shown, generic exemplary code shown in monospaced font. **c** Upper, shows this process for a small group of neurons from the telencephalon. Mirroring neurons to the same side produces cohesive clusters. Lower, neurons can be cut up (pruned) relative to user defined coordinates to, in this case, divide them by telencephalon region. NBLAST can be run separately on these two groups, and the results were compared using a tanglegram to see whether neurons that cluster together in the posterior telencephalon also cluster together in the ventral telencephalon.

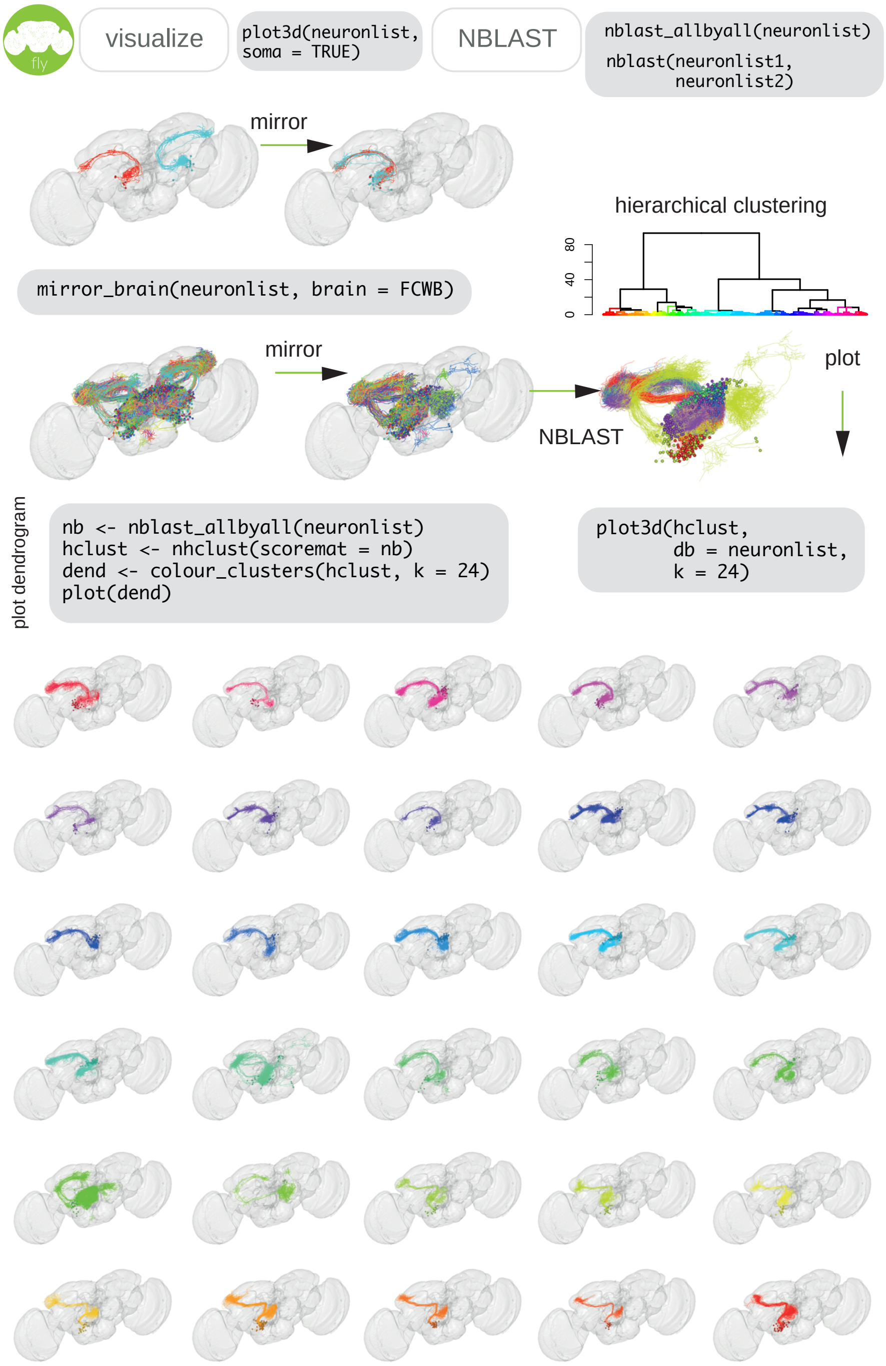


**Supplementary Figure 6:** The same basic process can be done for FlyCircuit neurons (Chiang et al., 2011), here subsetted (`in_volume`) to those neurons that have arbor in both the antennal lobe and lateral horn (i.e. olfactory projection neurons).

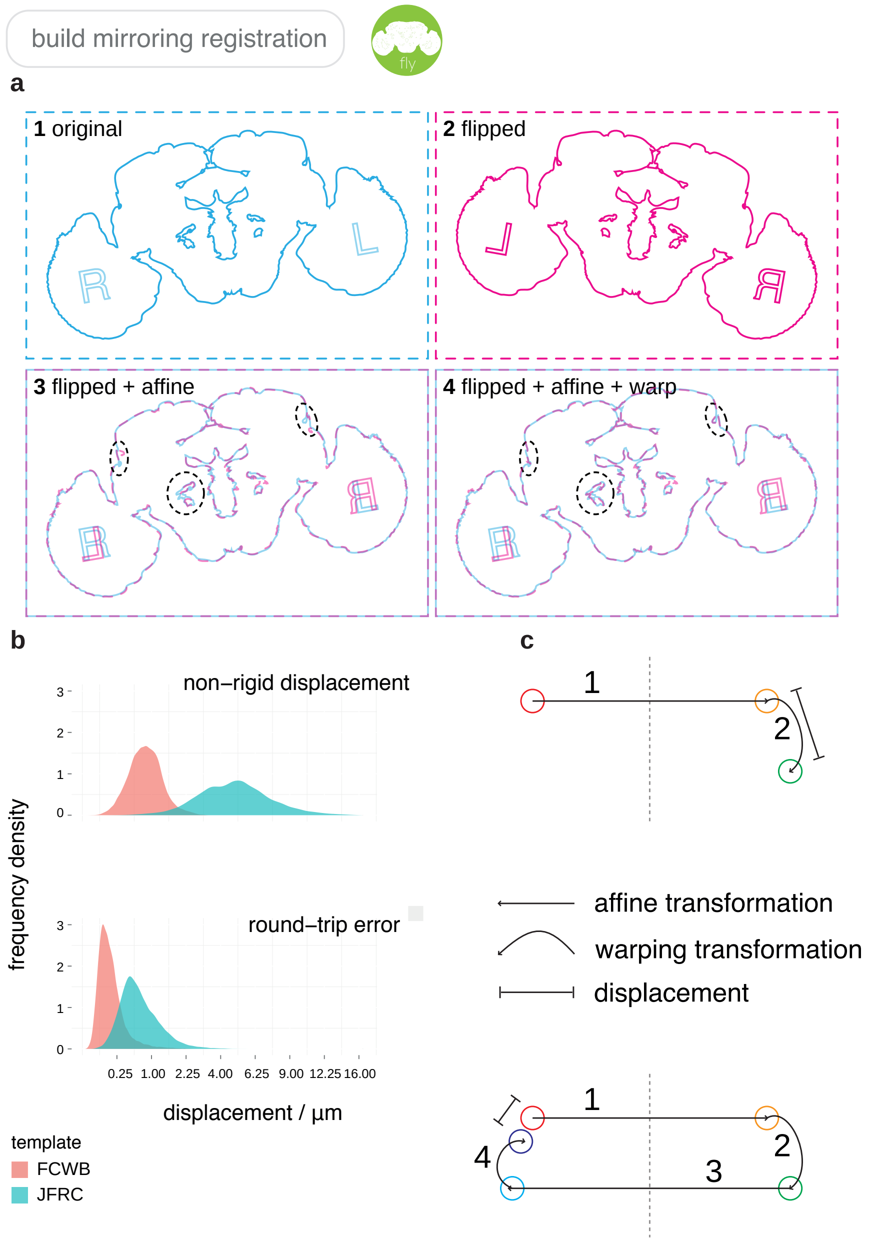


**Supplementary Figure 7: a** The full process undergone by an image during a mirroring registration. (1) Original image. (2) Flipped 180° around the medio-lateral axis. (3) Affinely transformed. (4) Non-rigidly warped. **b** Heatmaps of deformation magnitude fields for mirroring FlyCircuit and FlyLight reference brains. **c** Distribution of deformation displacements for both brains, in a single mirroring operation and a full round-trip. Illustrations show the transformations that points undergo.

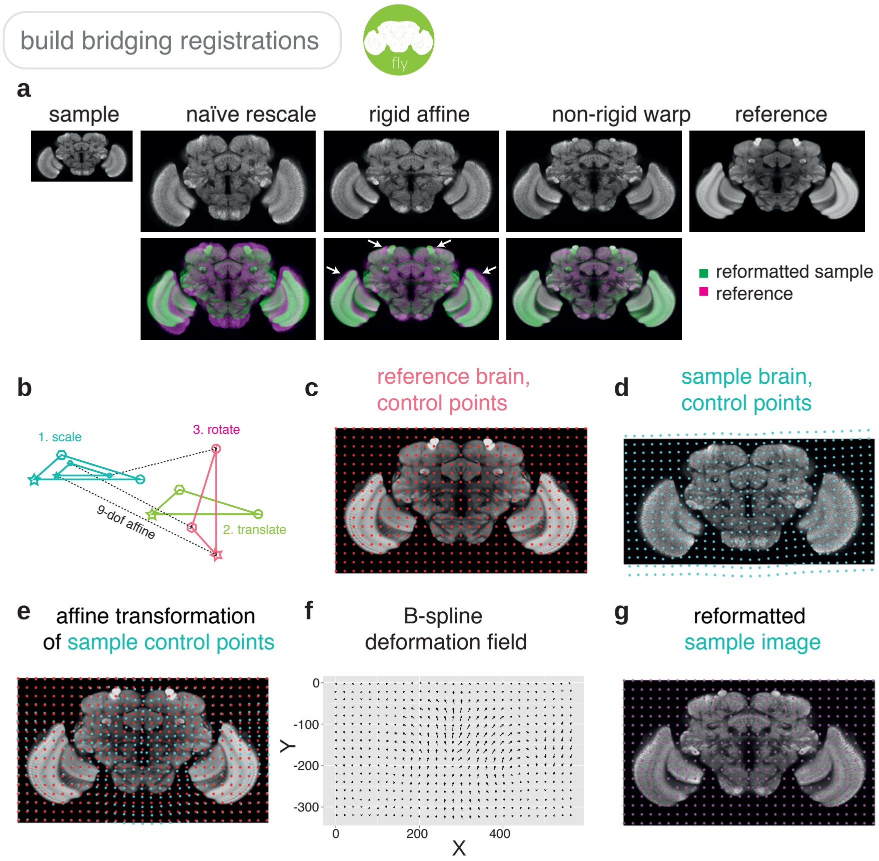


**Supplementary Figure 8: a** Increasing levels of registration complexity give increasingly good registration results. **b** Four composite transformations of a 12-degree-of-freedom affine transformation. **c** Regularly spaced grid of control points (red dots) in the reference brain. **d** Control points (cyan dots) in the sample image. **e** Sample brain control points affinely transformed into image space of reference brain, along with original control points. **f** Deformation field interpolated using B-splines. **g** Reformatted sample image.

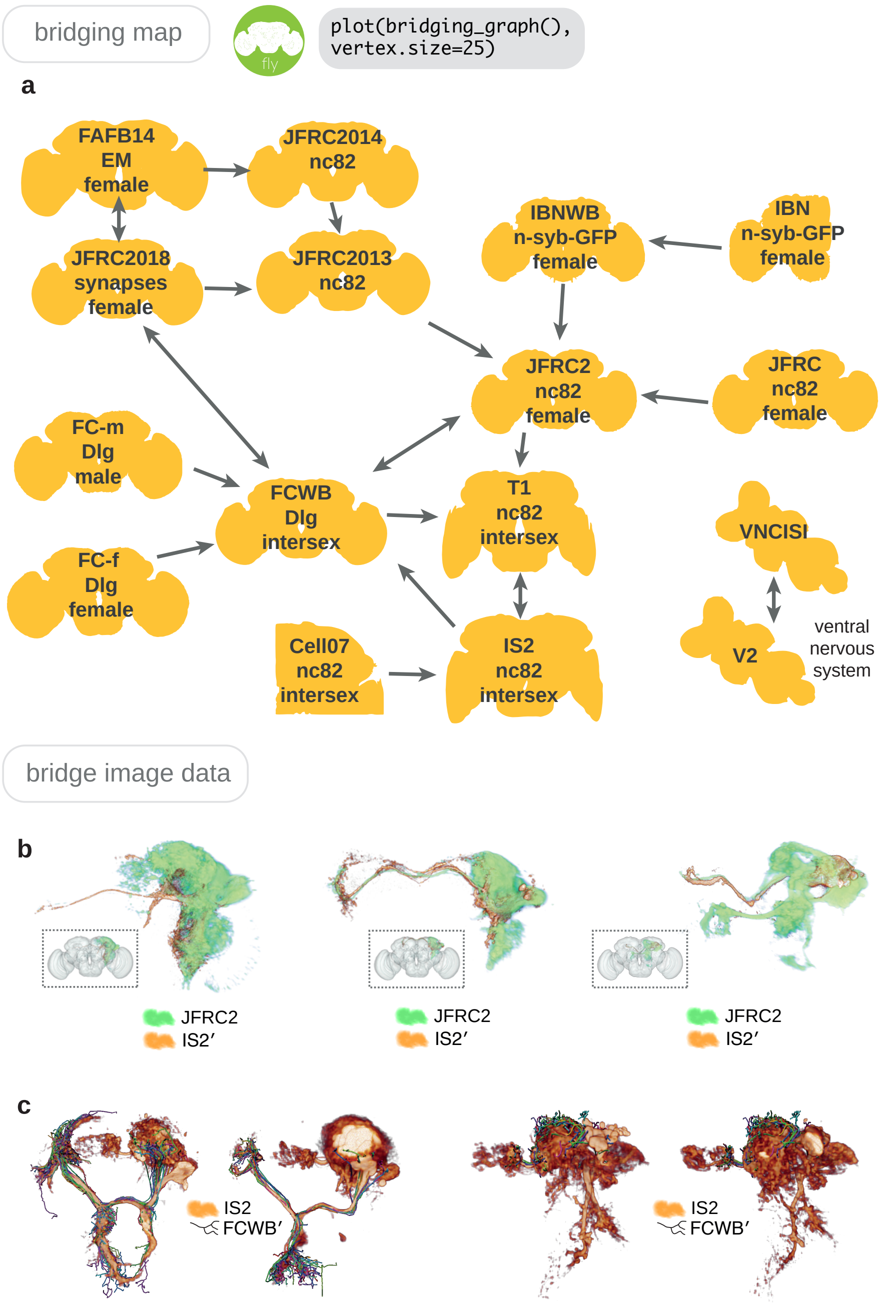


**Supplementary Figure 9: a** A sub-section of the bridging registrations available through nat.flybrains, via which a neuroanatomical entity in any brainspace can eventually be placed into any other brainspace, by chaining bridging registrations. Arrows indicate the existence and direction of a bridging registration. Registrations are numerically invertible and can be concatenated, meaning that the graph can be traversed in all directions. **b** Fru+ neuroblast clones (orange) transformed from IS2 space into JFRC2 space of elav neuroblast clones (green). **c** Sexually dimorphic Fru+ neuroblast clone (male on left, female on right) along with traced neurons from FlyCircuit.

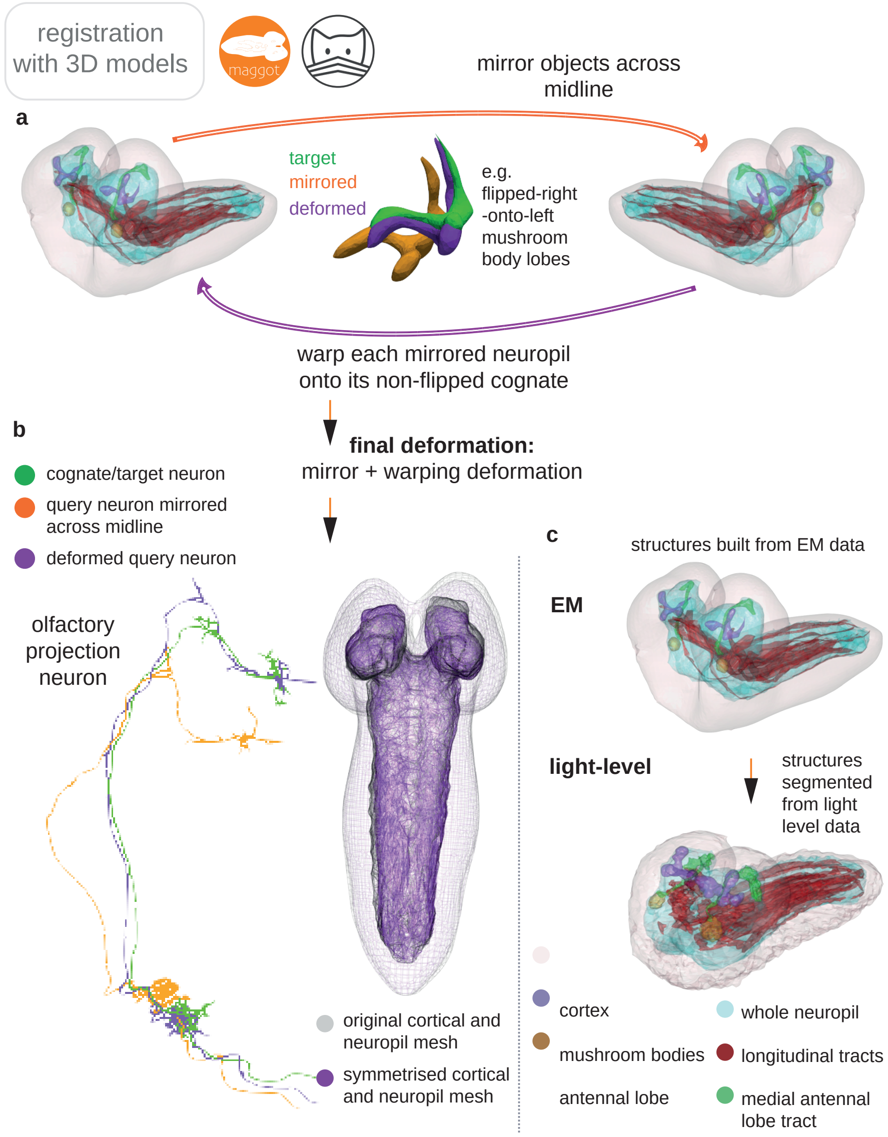


**Supplementary Figure 10: a** A left-right registration for the lop-sided nascent L1 larval connectome (Ohyama et al., 2015). Neuroanatomical models created from CATMAID data (upper) and 644 manually left-right paired neurons can be used as a basis to generate a single deformation of L1 space using Deformetrica (via deformetricar), that describes a left-right mapping in a single deformation of ambient space. **b** Symmetrisation (purple) of the lop-sided cortex and neuropil in the L1 EM dataset. **c** Registration of segmented light-level volumes onto the EM dataset. Immunohistochemical staining of dissected L1 central nervous system was needed to make 3D models for coarse neuroanatomical volumes, which could be matched to those in the L1 EM; namely, the pattern of longitudinal ventral nervous system axon tracts and the mushroom bodies, visible in a dFasciculin-II-GFP protein trap line, the laddered ventral nervous system pattern and olfactory tracts visible in a GH146-lexA line, and a Discs large-1/dN-Cadherin stain to visualize the neuropil and cortex of the brain.

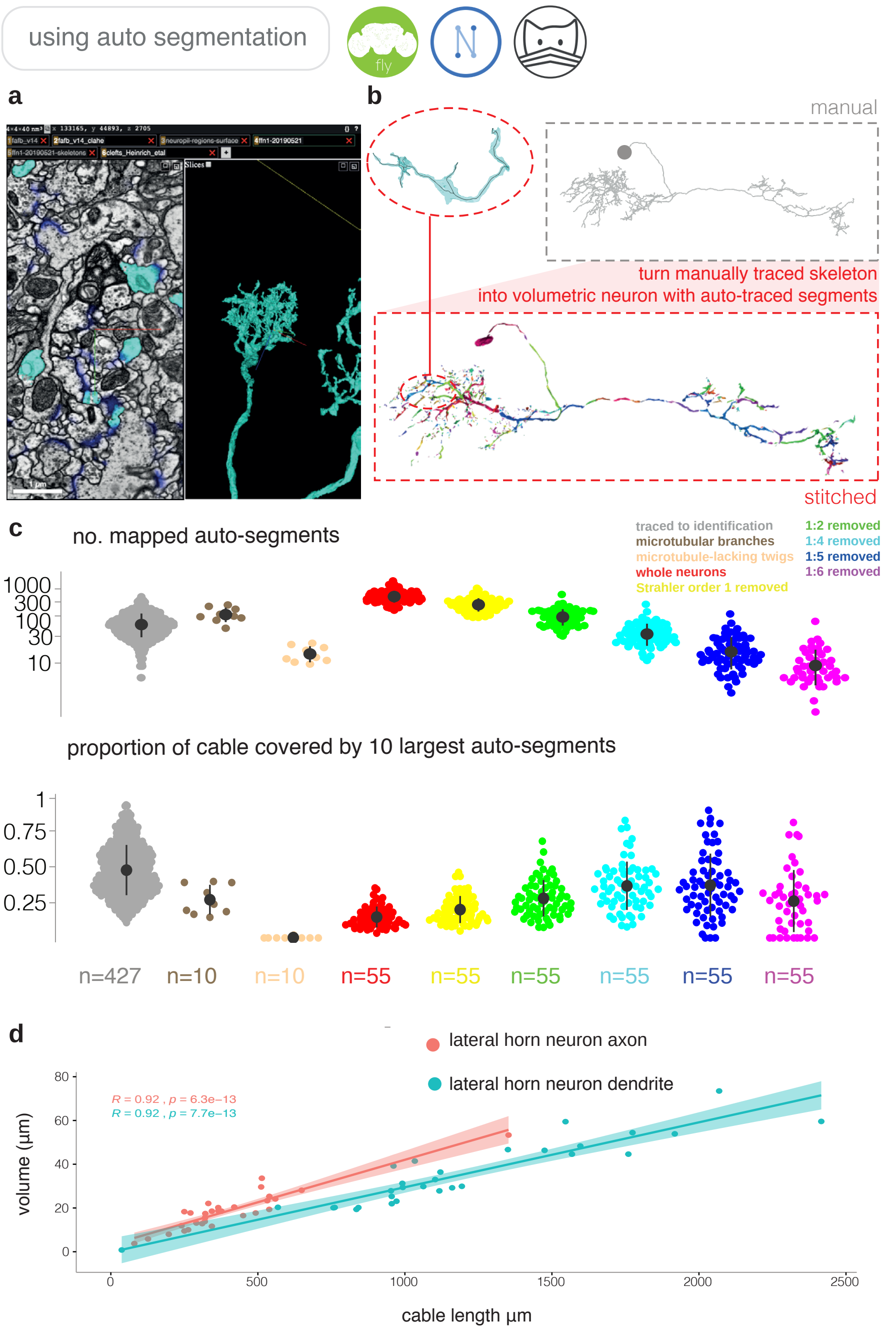


**Supplementary Figure 11: a** A NeuroGlancer window open on a web-browser, showing an example of an automatic reconstruction. **b** Automatically reconstructed segments can be mapped onto extant manual tracing in FAFB14 using our R package fafbseg, enabling easy volumentric reconstruction of neurons. **c** Upper, automatically traced segments are mapped onto 55 manually reconstructed neurons from the lateral horn of *D. melanogaster* in the FAFB14 dataset (Bates & Schlegel, in prep.), broken down by Strahler order, to simulate different levels of 'completeness'. A further 10 neurons have had microtubular cable annotated. Lower, proportion of cable at different levels of pruning, that are covered by the 10 largest auto-segmented fragments for each neuron. The traced to identification category comprises neurons reconstructed by expert annotators sufficiently for them to be identifiable in light level data, using the pipeline shown in Figure 8. **d** Correlation between cable length and volume for axons and dendrites (Schneider-Mizell et al., 2016) for a selection of central brain neurons.
